# NADP-dependent malic enzyme 1 participates in the abscisic acid response in *Arabidopsis thaliana*

**DOI:** 10.1101/348771

**Authors:** Cintia L. Arias, Tatiana Pavlovic, Giuliana Torcolese, Mariana B. Badia, Mauro Gismondi, Verónica G. Maurino, Carlos S. Andreo, María F. Drincovich, Mariel C. Gerrard Wheeler, Mariana Saigo

## Abstract

*Arabidopsis thaliana* possesses three cytosolic (NADP-ME1-3) and one plastidic (NADP-ME4) NADP-dependent malic enzymes. NADP-ME2 and-ME4 show constitutive expression, in contrast to NADP-ME1 and-ME3, which are restricted to particular tissues. Here, we show that NADP-ME1 transcript and protein were almost undetectable during normal vegetative growth, but gradually increased and reached levels higher than those of the other isoforms in the latest stages of seed development. Accordingly, in knockout *nadp-me1* mature seeds the total NADP-ME activity was significantly lower than in wild type mature seeds. The phenotypic analysis of *nadp-me1* plants indicated alterations of seed viability and germination. Besides, the treatment with abscisic acid (ABA), NaCl and mannitol specifically induced the accumulation of NADP-ME1 in seedlings. In line with this, *nadp-me1* plants show a weaker response of primary and lateral root length and stomatal opening to the presence of ABA.

The results suggest that NADP-ME1 plays a specialized role, linked to ABA signalling during the seed development as well as in the response to saline and osmotic stress.

## Introduction

NADP-dependent malic enzyme (NADP-ME; EC 1.1.1.40) catalyzes the oxidative decarboxylation of malate to generate pyruvate, CO_2_ and NADPH. In plants, the NADP-ME family is represented by several members, localized to cytosol and plastids. One of the better-established roles of this enzyme is the participation as malate decarboxylase in C4 and CAM photosynthesis (Drincovich *et al.*, 2011; Saigo *et al.*, 2013). Other functions are suggested based on the importance of malate balance for pH regulation, stomatal opening or lipogenesis (Laporte *et al.*, 2002; Hurth *et al.*, 2005; Gerrard Wheeler *et al.*, 2016). Arabidopsis possesses three cytosolic (NADP-ME1-3) and one plastidic NADP-ME isoforms (NADP-ME4; AT1G79750) (Gerrard Wheeler et al. 2005). NADP-ME2 (AT5G11670) is responsible for most of the NADP-ME activity measured in mature organs and has been involved in sugar metabolism in veins (Brown *et al.*, 2010) and in the oxidative burst triggered by hemibiotrophic fungal pathogen infection (Voll *et al.*, 2012). NADP-ME3 (AT5G25880) is only found in trichomes and pollen (Gerrard Wheeler et al. 2005).

Regarding NADP-ME1 (AT2G19900), its expression is very low in seedlings and adult plants, but significantly higher levels were found in maturing seeds and roots (Gerrard Wheeler *et al.*, 2005). This enzyme belongs to a particular phylogenetic group composed by NADP-MEs from different plant species (Gerrard Wheeler et al 2005). In *Zea mays*, a similar NADP-ME that is also specific of embryo roots, has shown similar expression pattern and kinetic characteristics such as lower catalytic efficiency and activation by succinate (Detarsio *et al.*, 2008; Alvarez *et al.*, 2013). Thus, the delimited localization and particular kinetic properties of NADP-ME1 probably reflect a particular role yet unknown. In a recent work based on transcriptome analysis and reverse genetics aimed at identifying differentially expressed genes during the imbibition and after-ripened seeds, *NADP-ME1* was found up-regulated in dormant Arabidopsis genotypes (Yazdanpanah *et al.*, 2017). The knock-out mutant *nadp-me1* showed disturbed seed traits compared to Col-0 plants (Yazdanpanah *et al.*, 2017).

Here, to disclose the biological role of *A. thaliana* NADP-ME1, we performed a deep analysis of its expression pattern measuring transcript accumulation, enzymatic activity and by using reporter genes. Besides, we evaluated the phenotypic parameters and physiological processes throughout Arabidopsis plant life affected by the absence of NADP-ME1. Overall, these findings indicate that there is a tight link of NADP-ME1 with processes related to abscisic acid (ABA) responses in seeds, roots and leaves.

## Materials and methods

### Plant lines, growing conditions and sampling

*Arabidopsis thaliana* Columbia-0 lines analyzed in this work include homozygous knockout mutants with T-DNA inserted into the genes encoding NADP-ME1 (*nadp-me1*; SALK_036898) and NADP-ME2 (*nadp-me2;* SALK_020607) and a triple mutant *nadp-me2×3×4* obtained by crosses (Gerrard Wheeler *et al.*, 2005). The position of the T-DNA insertion into each *NADP-ME* gene was verified by amplifying and sequencing the T-DNA flanking genomic DNA.

Arabidopsis transgenic lines were obtained by transforming wild type plants (Columbia-0, WT) with a construct carrying the complete coding sequence of NADP-ME1 fused to YFP (yellow fluorescent protein) gene, under the control of the *NADP-ME1* (referred as NADP-ME1::YFP) or the double 35SCaMV promoter. The construction that holds *NADP-ME1* promoter contains the 2,000 bp long sequence upstream transcription +1 site and the first intron of the gene. The binary vector ER-yb (Nelson *et al.*, 2007) was used. Inflorescences were incubated with cultures of *Agrobacterium tumefaciens* strain GV3101 using the protocol described in (Clough and Bent, 1998.) The transformed plants were selected with the herbicide BASTA. Four homozygous T3 lines for each construction were analyzed.

Seeds were sterilized with 0.5% (v/v) Triton X-100 and 50% (v/v) ethanol for 3 min, washed with 95% (v/v) ethanol and dried on filter paper. Seeds were stratified for 72 h at 277 K in the dark to synchronize germination, unless otherwise is stated. Plants were grown in 1X MS plates (Murashige and Skoog, 1962) or in soil in a culture room at days of 16 h of light with a flux density of 100 μE m^−2^ s^−1^ at 296-298 K. Seeds were collected at different stages, including 7, 12 and 18 days after pollination (DAP), mature (28 DAP), and 1 and 2 days after imbibition (DAI). Seedlings of 8 days grown in MS plates were transferred to plates supplemented with 100 mM NaCl, 225 mM mannitol or 0.5-10 μM ABA and collected at different times. All samples were frozen in liquid N_2_ and stored at 193 K.

### Real time polymerase chain reaction (qPCR) assays

Total RNA was extracted using a method developed for seed samples of Arabidopsis (Oñate-Sánchez and Vicente-Carbajosa, 2008) or a phenol-based one (Chomczynski and Sacchi, 1987) and plant RNA purification columns (PureLink, Amicon) for the rest of plant tissues, and then treated with RQ1DNase (Promega). The quantity and quality were evaluated by spectrophotometric measurements and electrophoresis in agarose gels. cDNAs were synthesized using MMLV (Promega) and random primers (Biodynamics). Relative expression was determined using specific primers (NADP-ME1left: 5-CAAGGCAATAAAACCGACTG-3´, NADP-ME 1 right: 5´-CATTTTTGCTAGTGGAAGCC-3´, NADP-ME2left: 5´-ACGATGGCAAAACCTACTTG-3´, NADP-ME2right: 5´-ATTGGCGTAATGCTCTTCTG-3´, NADP-ME3left: 5´-GGCACCAATCAGACTCAGATCT-3´, NADP-ME3 right: 5´-AGCAAGTCCTTTATTGTAACGT-3´, NADP-ME4left: 5´-CTTTCGAACCCAACTTCTCA-3´, NADP-ME4right: 5´-CATTATTAGCCCGAGTCCAA-3´, YFPleft: 5´-ACGTAAACGGCCACAAGTTC-3´, YFPright: 5´-AAGTCGTGCTGCTTCATGTG-3´) and polyubiquitin 10 gene(AT4G05320; Czechowski *et al.*, 2005) as normalizer. The amplifications were performed on a Stratagene Mx3000P cycler, using the SYBR Green I dye (Invitrogen) as a fluorescent reporter. PCR controls were made to ensure that the RNA samples were free of DNA contamination. The PCR specificity was verified by melting curve and gel electrophoresis analysis of the products. The relative expression was calculated using a modified version of the 2^−ΔΔCt^ method (Pfaffl, 2001), the efficiencies and the propagation of errors determined according to (Liu and Saint, 2002) and (Hellemans *et al.*, 2007).

### Microscopy analysis

Detection of YFP was achieved using a YFP filter (excitation, 488 nm; emission, 505-550 nm) and a Karl Zeiss Lsm880 or a Nikon Eclipse TE-2000 Model-E2 confocal microscope. Roots were mounted in propidium iodide dye (Invitrogen) and the imaging settings were 488 nm excitation and >585 nm emission.

### NADP-ME activity measurements in extracts and western blot analysis

Samples were homogenized in mortars according to (Badia *et al.*, 2015) and the extracts were desalted through Sephadex G-50 spin columns. Protein concentration was determined by the BioRad protein assay using total serum protein as standard. NADP-ME activity was assayed at 303 K in a Jasco spectrophotometer following the appearance of NADPH at 340 nm (ε_340nm_=6.22 mM^−1^ cm^−1^) using 50 mM MOPS-KOH pH 6.8, 10 mM MgCl_2_, 0.5 mM NADP and 10 mM malate. One unit (U) is defined as the amount of enzyme that catalyzes the formation of 1 μmol of NADPH min^−1^ under the specified conditions.

SDS-PAGE was performed in 10% (w/v) polyacrylamide gels according to Laemmli, 1970. Proteins were then electroblotted onto a nitrocellulose membrane. Antibodies against green fluorescent protein (Abcam), which also immunodetect YFP fusion proteins, were used. Bound antibodies were visualized by linking to alkaline phosphatase conjugated goat anti-rabbit IgG according to the instructions of the manufacturer (Sigma). Alkaline phosphatase activity was detected colorimetrically.

### Determination of phenotypic parameters

The analyzed lines were grown simultaneously with a randomized physical arrangement and frequently rotated. Germination was evaluated by counting the number of seeds with visible radicles, in plate growth assays without previous stratification. Seeds freshly collected or stored at room temperature for 1-11 years were used. For primary and lateral root length measurement, 5-day-old seedlings were transplanted to MS plates without or with 10 μM ABA and monitored for 6 days.

### Controlled deterioration test

The controlled deterioration test was performed as described previously (Mao and Sun, 2015) with minor modifications. Briefly, freshly harvested seeds were dried in a desiccator containing silica gel and then equilibrated for 3 days at 288 K and 85-90% relative humidity (RH) in a hygrostat of KCl. Then, the hygrostat was transferred to 313 K which resulted in 80-85% RH. After 1-7 days at high temperature, the seeds were stored at 293 K and 33% RH for 3 days in a hygrostat of MgCl_2_ and dried again in a desiccator with silica gel (6% RH). The RH and temperature were monitored in all steps with a datalogger. The germination was assayed in replicates of 100 seeds in agar plates and recorded after 7 days.

### Stomatal opening assays

WT and *nadp-me1* plants were grown in soil pots for 2-3 weeks. The stomatal response was evaluated in detached leaves measuring the stomatal aperture after treatment with 30 μM ABA according to (Wohlbach *et al.*, 2008).

### *In silico* phylogenetic analysis

Protein sequences were retrieved from Phytozome 9.1 database (www.phytozome.net), using *A. thaliana* NADP-ME2 as query. The evolutionary history was inferred using the Neighbor-Joining method (Saitou and Nei, 1987). The evolutionary distances were computed using the Poisson correction method (Zuckerkandl and Pauling, 1965). The analysis involved 40 amino acid sequences. All positions containing gaps and missing data were eliminated. There were a total of 488 positions in the final dataset. Evolutionary analyses were conducted in MEGA7 (Kumar *et al.*, 2016). The detection of ABA related elements was performed with a phylogenetic footprinting tool with a set of characterized motifs (Table S1, cis-analyzer, Gismondi unpublished).

### Statistical analysis

Significance was determined by the ANOVA or Student’s t test using the SigmaPlot software. For the case of qualitative variables, such as germination analysis, the data were analyzed using comparisons of proportions through the Z statistic. The sample size is indicated in each figure or table.

## Results

### *NADP-ME1* increases during seed maturation in Arabidopsis

The expression profile of NADP-ME1 was analyzed along Arabidopsis seed maturation and germination. The transcript level of *NADP-ME1* sharply increases during seed maturation from 7 DAP until complete maturation, reaching a value 136-fold higher at 18 DAP (Fig. 1A). By contrast, the transcript levels of *NADP-ME2*, *NADP-ME3* and *NADP-ME4* remain fairly constant along seed maturation (Fig. 1A). In germinating seeds the transcript level of *NADP-ME1* drastically decreases to the low levels detected at 7 DAP (Fig. 1A). A similar pattern of *NADP-ME1* expression was observed using the YFP reporter gene under the control of *NADP-ME1* promoter. In embryos, YFP fluorescence is undetectable up to 9 DAP; increase from 11 to 18 DAP; and then decrease in germinating embryos (Fig. 1D).

**Fig. 1:**
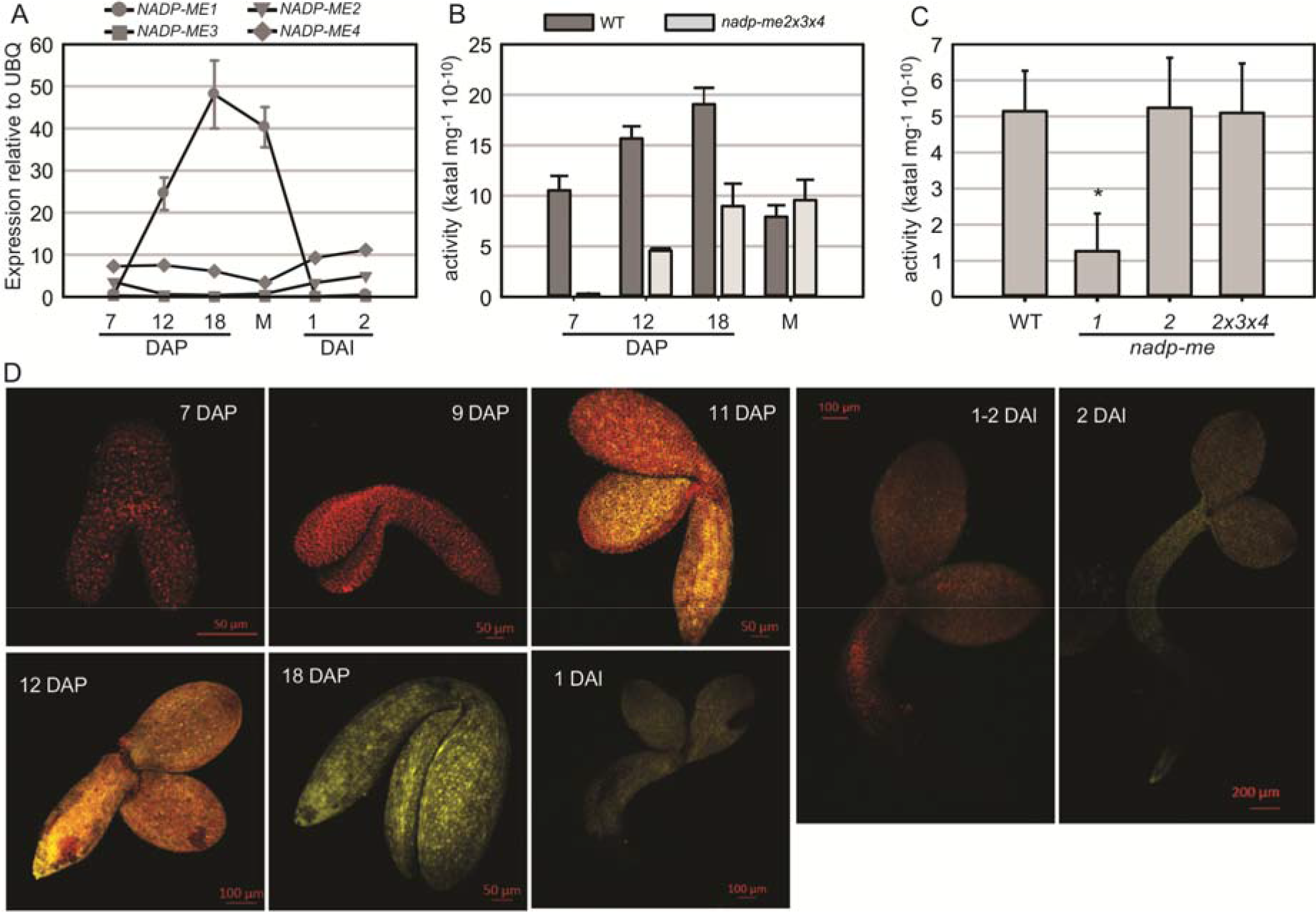
NADP-ME1 expression in Arabidopsis seeds. (A) Relative level of the transcripts of *NADP-ME* genes in seeds collected at 7, 12 and 18 days after pollination (DAP), mature stage (M), and at 1 and 2 days after imbibition (DAI). The polyubiquitin 10 (UBQ) gene was used as reference. NADP-ME activity was assayed in siliques throughout maturation (B) and in mature seeds (C) for WT and simple or triple mutant lines. The values are the average of at least two independent experiments ± SD. Values marked with an asterisk (*) indicate that are significant different (p < 0.05). (D) Expression of the fusion protein NADP-ME1::YFP during seed development and germination. The reporter contains the complete coding sequence of NADP-ME1 fused to YFP gene, under the control of the *NADP-ME1* promoter. YFP fluorescence in representative embryos at 7; 9; 11; 12 and18 DAP and at 1, 1-2 and 2 DAI are shown. Scale bars are indicated in each panel.

NADP-ME activity in Arabidopsis siliques increases from 7 DAP to 18 DAP (Fig. 1B), following a similar profile as that observed for *NADP-ME1* transcript level (Fig. 1A). The increase of NADP-ME activity in *nadp-me2×3×4* triple mutant, in which NADP-ME1 is the only NADP-ME found, matches the increase of activity found in WT. Thus, the NADP-ME activity profile in Arabidopsis WT may be endorsed to NADP-ME1 increase (Fig. 1B). When comparing NADP-ME activity in mature seeds of WT and *nadp-me* mutant lines, the lack of NADP-ME2 alone or in combination with NADP-ME3 and NADP-ME4 does not affect the total NADP-ME activity. In contrast, a drastic decrease of NADP-ME activity is observed in *nadp-me1* knockout mutant (Fig. 1C). Overall, it is clear that NADP-ME1 is the isoform that contributes the most to NADP-ME activity at mature seed stage (Fig. 1C).

### *NADP-ME1* is up-regulated by NaCl, mannitol and ABA in Arabidopsis seedlings and roots

*NADP-ME1-4* transcript levels were assayed in Arabidopsis rosettes of 8 days (Fig. 2A, Supplementary fig. 1A). As previously shown (Gerrard Wheeler et al 2005), the transcripts of *NADP-ME1* and *−3* display very low levels in relation to *NADP-ME2* and *−4* in control conditions (MS, Supplementary fig. 1A). However, when 8-day-old seedlings are treated with 100 mM NaCl; 225 mM mannitol or 10 μM ABA, a strong induction *NADP-ME1* transcript is observed (53, 79 and 55 times, respectively). Neither *NADP-ME2* nor *NADP-ME3* or *NADP-ME4* shows such a significant response as *NADP-ME1* (Fig. 2A, Supplementary fig. 1A). We further analyzed the response of 8-day-old seedlings to different ABA concentrations (0.5, 1, 5 and 10 μM of ABA) and exposure times. The level of *NADP-ME1* transcript increases 5-6 folds in 0.5, 1 and 5 μM ABA and 10 folds in 10 μM ABA, reaching a level which is almost twice as high as that of *NADP-ME2* (Supplementary fig. 1B). When the length of the ABA treatment was tested, we found that *NADP-ME1* increases 6 and 12 folds at 6 and 12 h, respectively and *NADP-ME2* levels did not varied significantly (Supplementary fig. 1C).

**Fig. 2:**
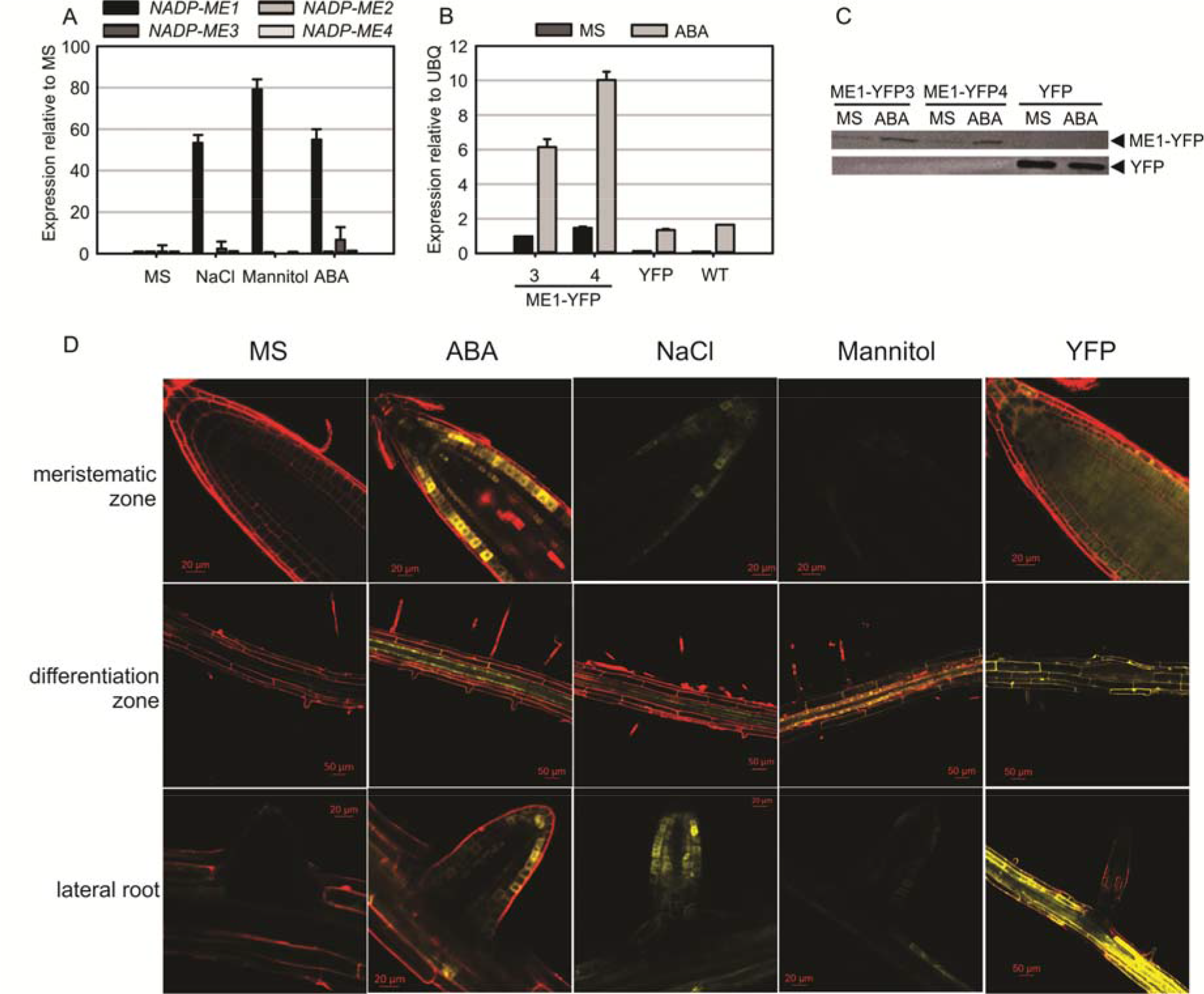
NADP-ME1 expression in response to NaCl, mannitol and ABA. (A) Relative levels of the transcripts of *NADP-ME* genes in rosettes in control conditions (MS) or after 6 hs treatments of seedling with 100 mM NaCl, 225 mM mannitol or 10 μM ABA. The polyubiquitin 10 gene (UBQ) was used as reference. The values are the average of at least two independent experiments ± SD. (B) Response of *NADP-ME1* promoter in transgenic lines. ME1-YFP3 and 4 are two independent transgenic lines expressing NADP-ME1::YFP under the control of *NADP-ME1* promoter. YFP denotes a line expressing the YFP coding sequence under the control of the double 35SCaMV promoter. The level of NADP-ME1 was compared in control conditions (MS) and after treatment with 10 μM ABA during 6 h was applied. (C) Western blot of the seedling protein extract (30 μg) from the transgenic lines. Molecular mass markers were run in parallel and stained with Coomassie Blue to localize the position of fusion and YFP proteins. (D) Expression of the fusion protein NADP-ME1::YFP in roots in response to ABA, NaCl and mannitol. YFP fluorescence in different parts of roots of 8-day-old seedlings incubated with 10 μM ABA, 100 mM NaCl or 225 mM mannitol for 6 h. Scale bars are indicated in each panel. Right panels show the fluorescence distribution of the control YFP line.

To test if NADP-ME1 protein was affected by the presence of ABA, we used the transgenic NADP-ME1::YFP lines to immunodetect the fusion protein. Consistent with the previous observations, a strong NADP-ME1 induction by ABA at the level of transcript and protein was observed in the lines expressing NADP-ME1::YFP under the control of *NADP-ME1* promoter (Fig. 2 B and C).

To further analyze the expression pattern of NADP-ME1 under water stress and ABA treatments we observed the YFP fluorescence in roots of 8-day-old seedlings of the transgenic lines expressing NADP-ME1::YFP under the control of *NADP-ME1* promoter. ABA, NaCl and mannitol produced similar induction of the NADP-ME1, especially in the root apical meristem, the differentiation zone and in lateral roots (Fig. 2D).

### Seeds of *nadp-me1* mutant are less sensitive to the ABA repression of germination and loss viability more rapidly than WT

The number of *nadp-me1* and WT germinated seeds with visible radicles was counted at different times after seeding. Almost 100% of *nadp-me1* seeds germinate approximately 50 hours after seeding in MS medium but WT seeds reach that value almost 48 hours later (Fig. 3A). In the presence of exogenous ABA, the percentages of germinated seeds are lower for both lines but *nadp-me1* seeds reach 2.5-3 folds higher germination percentages than WT at 62 and 86 hours (Fig. 3B). Then, these results show that under both conditions the germination rate of *nadp-me1* is faster than WT and a lower sensitivity of *nadp-me1* seeds to ABA repression of germination.

**Fig. 3:**
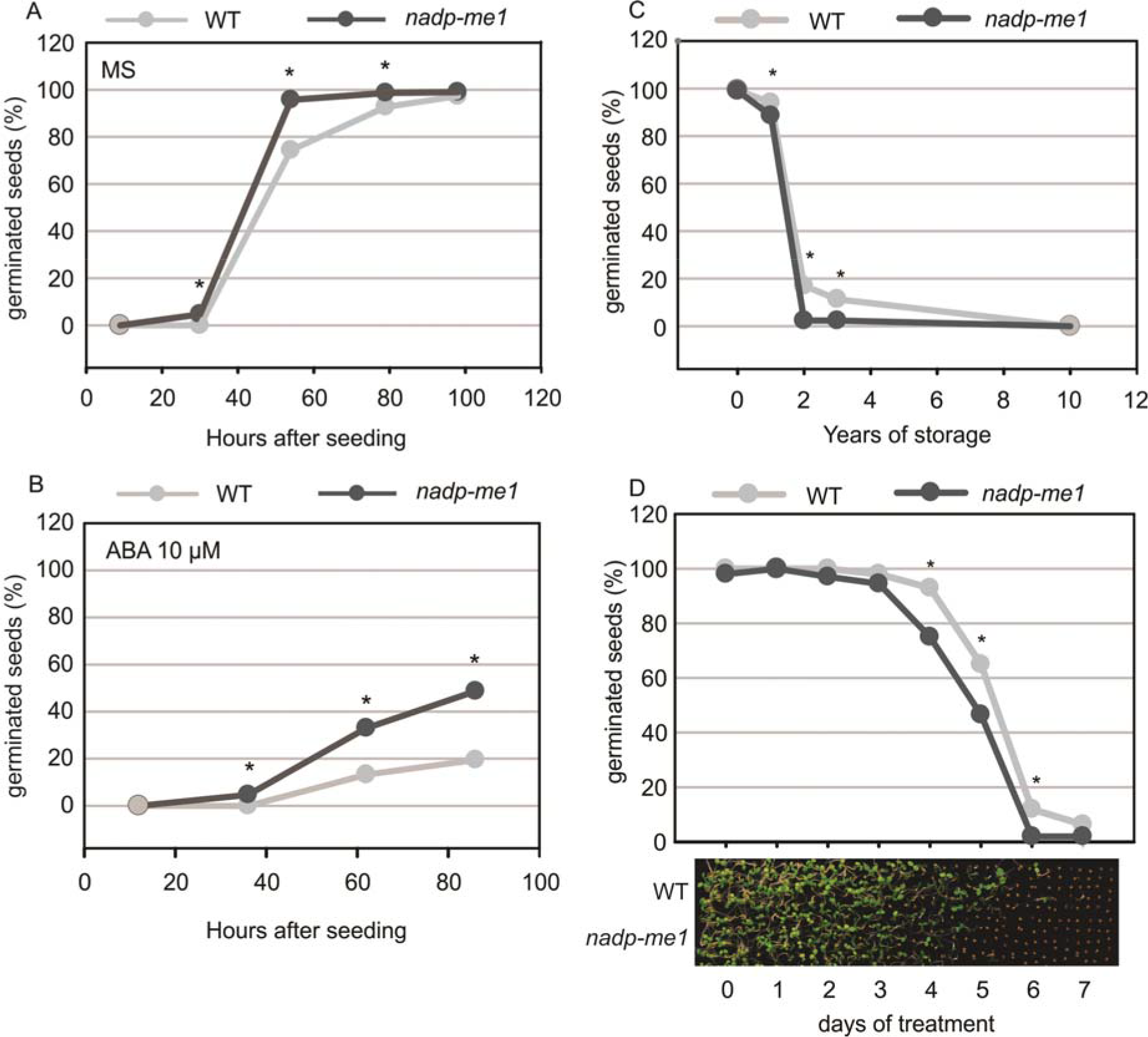
Germination and seed viability assay in *nadp-mel* mutant. Germination was evaluated by counting the number of seeds with visible radicles in MS plates (A) or MS supplemented with 10 μM ABA (B). The proportion of germinated seeds after evaluating 300 seeds per line is shown. Germination was also evaluated using long time stored seeds (C) or after exposing them to high temperature (313 K) for 1-7 days (D). The asterisk denotes significant differences (p < 0.05) between WT and *nadp-me1* lines.

Seed viability was also analyzed for the *nadp-me1* mutant. The viability of recently harvested *nadp-me1* and WT seeds is 100%. However, when we tested seeds stored for long times, we found that *nadp-me1* seeds loss viability earlier than WT (Fig. 3C).

To examine whether the longevity is affected in seeds lacking NADP-ME1, we performed a controlled deterioration test based on the exposure of seeds to high temperature (40°C) and relative humidity (80-85%) for several days. Although the viability is the same as WT without treatment and after complete treatment (7 days), *nadp-me1* seed decay is faster than WT (Fig. 3D).

### Stomata and roots in knockout *nadp-me1* are less sensitive to ABA

Accordingly to the very low levels of *NADP-ME1* expression observed in leaves, *nadp-me1* mutant plants do not show differences in stomatal aperture compared to WT under normal conditions. However, in the presence of 30 μM ABA the stomata pore size is larger in plants lacking NADP-ME1 than in WT (Fig. 4A).

**Fig. 4:**
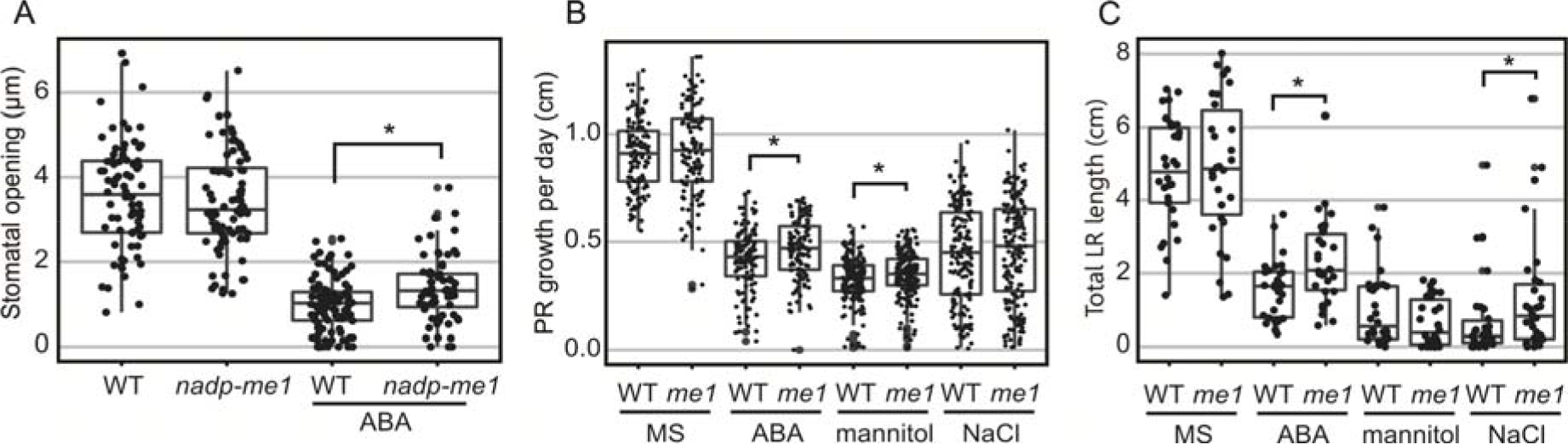
Stomatal opening and root length assays in *nadp-me1* mutant. (A) Stomata pore size was determined in the light and in the presence or absence of 30 μM ABA. Between 70-100 stomata aperture were measured. Primary (PR, B) and lateral (LR, C) root length were assayed in seedling transferred to MS plates supplemented with 10 μM ABA, 100 mM NaCl or 225 mM mannitol for 6 days. All values are presented and the statistical descriptions of each set of data are shown as box plots. The asterisk denotes significant differences (p < 0.05) between WT and *nadp-me1* line.

Root growth responses to ABA, mannitol and NaCl are also different in *nadp-me1* and WT. When 5-day-old seedlings are transferred to MS medium supplemented with 10 μM ABA, increased primary root (PR) elongation rate and total length of lateral roots (LR) are found in *nadp-me1* mutants with respect to WT (Fig. 3B and C). When the seedlings are transferred to MS supplemented with 100 mM NaCl, the rate of growth of the PR is not significantly different for *nadp-me1* mutants compared to WT, but the average total length of LR is higher for the mutant plants (Fig. 3B and C). On contrary, mannitol treatment inhibits the PR growth to a lesser extent in *nadp-me1* plants compared to WT and do not show a significant effect on the total length of LR (Fig. 3B and C).

## Discussion

### NADP-ME1 expression is up-regulated by ABA and its absence weakens the ABA response in different Arabidopsis organs

Despite *A. thaliana* having three cytosolic NADP-ME isoforms, NADP-ME1 displays a distinctive and specific expression pattern. Our results indicate that under normal conditions, *NADP-ME1* transcript is detected almost exclusively in maturing seeds (Fig. 1), while it accumulates in the rosettes and roots under saline or osmotic stresses (Fig. 2). Besides, we found that ABA increases *NADP-ME1* transcript, which is correlated with an increase in NADP-ME1 protein (Fig. 2A B and C). This up-regulation is exclusively exerted over *NADP-ME1*, since *NADP-ME2*, *NADP-ME 3* and *NADP-ME 4* do not show this response in the conditions assayed here.

Considering that ABA mediates part of the response to saline and osmotic stress and the regulation of NADP-ME1 expression by ABA treatment, the sharp increase in the expression of *NADP-ME1* in maturing seeds could be caused by the accumulation of ABA (Fig. 1A). Thus, it seems that this phytohormone acts as one of the major signals controlling *NADP-ME1* expression.

Besides the control of NADP-ME1 expression exerted by ABA, we show that the absence of NADP-ME1 affects ABA response in different Arabidopsis organs. Particularly, we found that the lack of NADP-ME1 affects not only the longevity of the seeds but also the control of the germination (Fig. 3). *nadp-me1* mutants exhibit less tolerance to prolonged storage and are less sensitive to the inhibition of the germination exerted by ABA (Fig. 3). Plants lacking NADP-ME1 also exhibit less sensitivity than the WT to stomatal closure and root growth inhibition induced by ABA, NaCl and mannitol (Fig. 4A and B).

It is well-known that ABA regulates vital processes associated to normal late seed development such as synthesis of reserve compounds, tolerance to desiccation, dormancy, longevity, and germination (North *et al.*, 2010). In addition, this phytohormone accumulates in large quantities during saline, osmotic or drought stress conditions and controls root growth and transpiration through stomatal closure (Priest *et al.*, 2006; Christmann *et al.*, 2007). ABA target genes belong to functional categories such as seed maturation (oleosins, dehydrins, or late embryogenesis abundant proteins), protein stability (proteases), cellular structure (expansins and wall synthesis enzymes), signalling (kinases and phosphatases), response to stress (heat shock proteins) and metabolism (Reeves *et al.*, 2011). Here, we found that there is a link between NADP-ME1 expression and ABA in *Arabidopsis thaliana*. Moreover, our results suggest that NADP-ME1 would be important to cope with conditions of water deficit, where the plant responses are mainly mediated by ABA.

### NADP-ME1: an old and conserved NADP-ME with particular physiological roles linked to ABA response

The sequence analysis of the complete set of NADP-ME isoforms from the dicot species *A. thaliana*, *Glycine max*, *Medicago truncatula*, *Phaseolus vulgaris*, *Ricinus communis* and the monocot species *Oryza sativa*, *Setaria italica* and *Zea mays* showed that monocot cyt2, cyt3 and plastidic isoforms belong to a monophyletic cluster independent from dicot cyt2 and plastidic isoforms that also cluster together (Fig. 5A). On the other hand, monocot cyt1 and dicot cyt1 groups show origins different from the rest of the family members and are more similar to the ancestral NADP-ME present in the origin of angiosperms (Fig. 5A). In different studies it has been shown that many of these *NADP-ME* cyt1 genes have expression patterns similar to *NADP-ME1*. In *G. max* and *R. communis* the transcripts encoding the isoforms homologous to NADP-ME1 also showed an induction along the seed maturation (Gerrard Wheeler *et al.*, 2016). The study of the *Z. mays* NADP-ME family showed that one cytosolic isoform also accumulated along the grain maturation and increased after ABA treatment in roots (Detarsio *et al.*, 2008; Alvarez *et al.*, 2013). Furthermore, Chi and collaborators (2004) observed that a cytosolic isoform of *O. sativa* is specifically induced by mannitol and NaCl and confers salt tolerance in transgenic overexpressing Arabidopsis (Chen and Long, 2007). These results suggest a conserved role for cyt1 NADP-ME proteins both in monocot and dicot species.

**Fig. 5:**
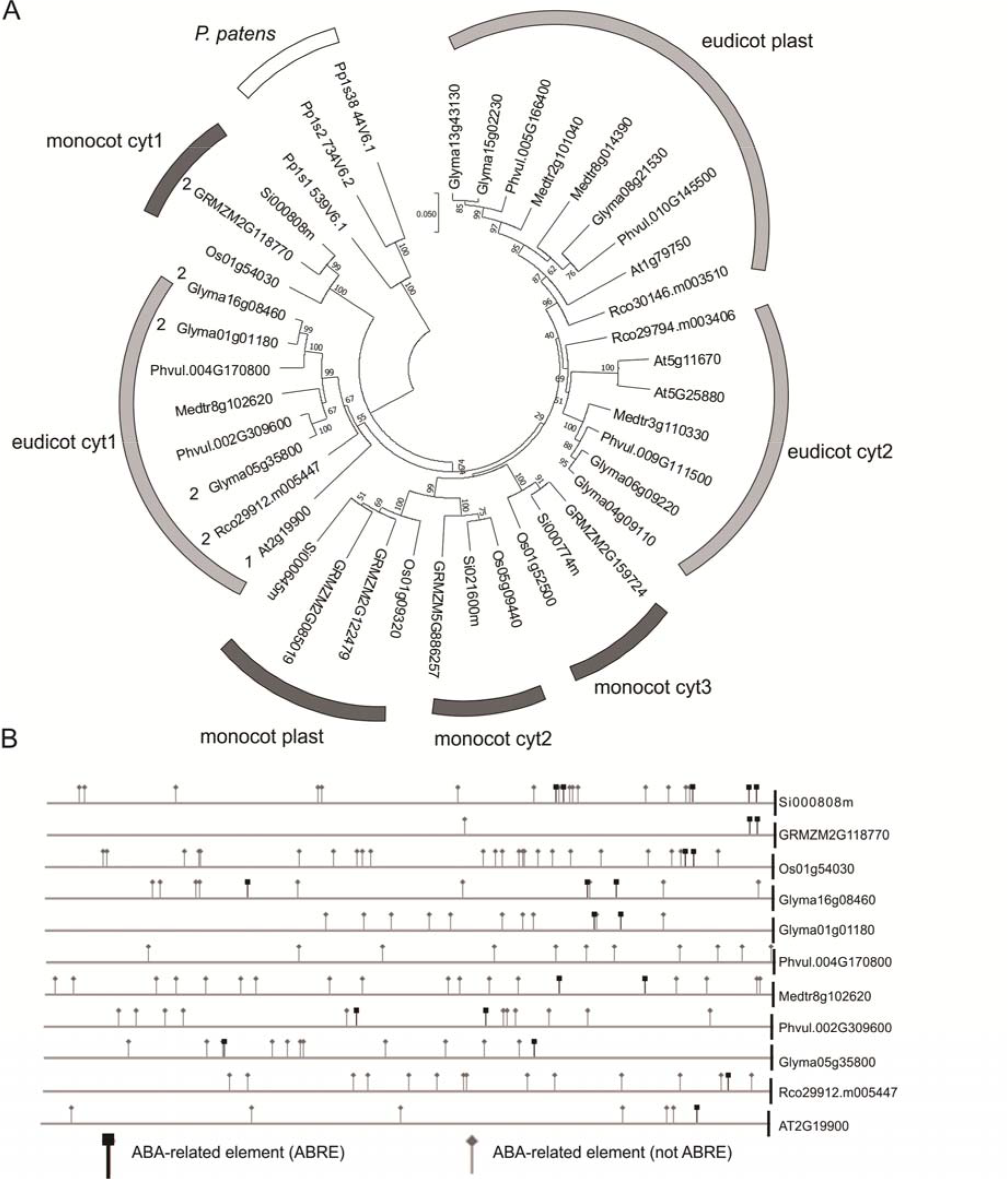
ABA and drought response elements in the promoters of *NADP-ME1* and related genes. (A) The evolutionary history of the complete set of NADP-ME isoforms from the dicot species Arabidopsis (At), soybean (Glyma), *Medicago truncatula* (Medtr), *Phaseolus vulgaris* (Phvul), *Ricinus communis* (Rco) and the monocot species rice (Os), *Setaria italica* (Si) and maize (GRMZM) was inferred using the Neighbor-Joining method. The optimal tree with the sum of branch length=2.44632768 is shown. The tree is drawn to scale, with branch lengths in the same units as those of the evolutionary distances used to infer the phylogenetic tree. The evolutionary distances are in the units of the number of amino acid substitutions per site. (B) The occurrence of ABA responsive elements was analyzed in the promoters of monocot cyt 1 and dicot cyt1 groups.

*In silico* analysis of the promoter and 5’UTR region of monocot cyt1 and dicot cyt1 genes showed that they conserve elements that are linked to the ABA response (Fig. 5B, Tables S1 and S2). In this sense, we observed an increment of the *NADP-ME1* transcript level in response to dose and time of ABA treatment (Supplementary Fig. 1B and C). It is also important to mention that these regulatory elements are also present in the bryophyte *Physcomitrella patens*, suggesting an ancestral and strongly conserved regulation of *NADP-ME* genes (Fig. 5 and Table S3). In spite of the evolutionary distance of this bryophyte with respect to angiosperms, ABA is also involved in tolerance to stress by balancing the water level in its tissues (Takezawa *et al.*, 2011). This could indicate that the participation of NADP-ME in the response to ABA existed in plants even before the appearance of stomata, vascular system and seeds.

Based on the phylogenetic and promoter analysis and in the phenotypic analysis carried out here, we propose that NADP-ME1 is an isoform that plays a very specialized role compared to the rest of the NADP-ME family members, possibly linked to the signaling initiated by ABA, as an intermediary or final effector (Fig. 6). This enzyme could fulfill its role by consuming malate and/or generating pyruvate and NADPH in the cytosol. Malate is a metabolic intermediate that is being recognized for its regulatory functions (Finkemeyer *et al.*, 2013). For example, the decrease of ion entrance through a vacuolar channel that promotes the stomata closure is regulated by the decrease of the cytosolic malate concentration (De Angeli *et al.*, 2013). Besides, the growth and the response of the roots to the availability of water also depend on the hydric potential generated by the movement of osmolytes through cellular compartments (Robbins and Dinneny, 2015). On the other hand, the NADPH generated could be used for the biosynthesis of compounds that accompany the stress response or to control reactive oxygen species generation. Although future experiments are necessary to elucidate the mechanism by which NADP-ME1 participates in the ABA signaling pathway, the results presented here clearly show the participation of this particular isoenzyme from the NADP-ME family in the response to a hormone which is a key in different physiological responses in plants. NADP-ME1 could work as an intermediate or end effector, possibly regulating the concentration of the reaction substrates and/or products to feed metabolic pathways and regulatory functions triggered by the water stress (Fig. 6). The findings shown here contribute to the understanding of the maturation and germination seed processes and the response of plants to water stress, and may help us to establish innovative strategies to generate crops with a better use of natural resources.

**Fig. 6:**
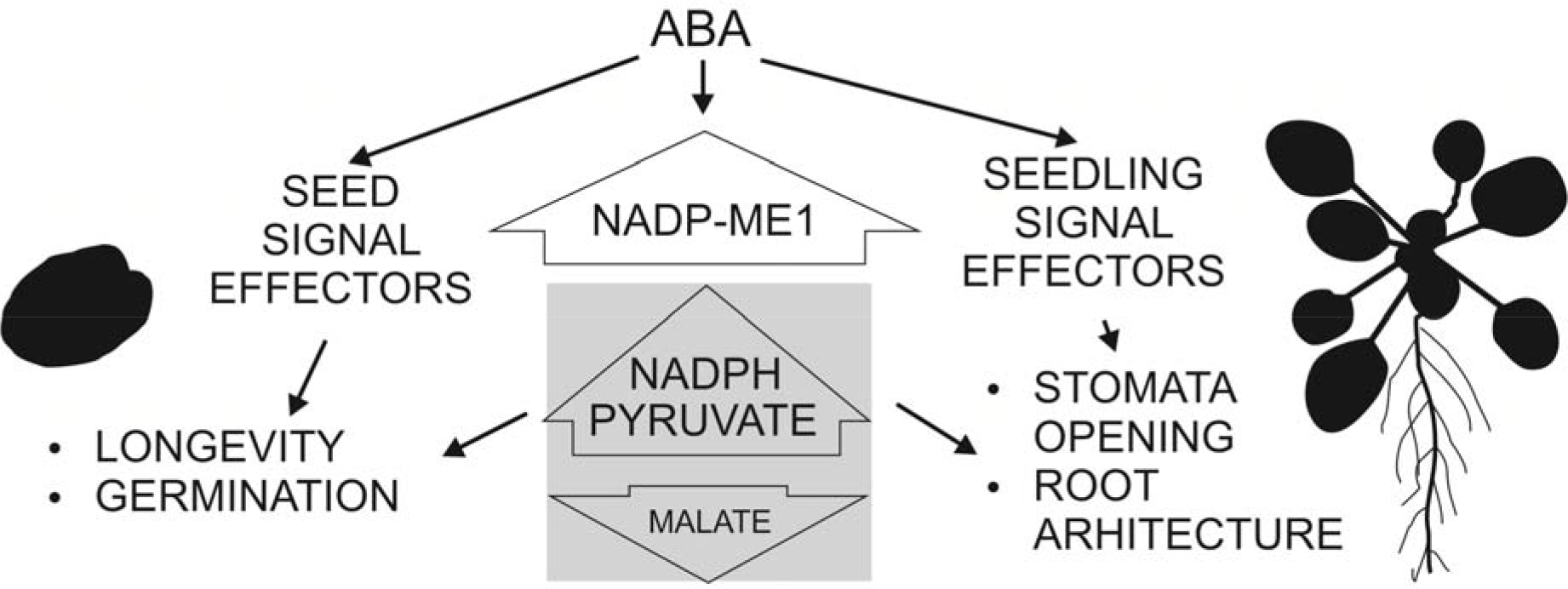
Schematic scheme showing the participation of NADP-ME1 in ABA response in Arabidopsis. ABA induces NADP-ME1 expression in particular organs and cell types producing a modification of malate, pyruvate and NADPH levels, which impact on seed longevity and germination, stomata opening and root architecture.

Abbreviations: ABA, abscisic acid; DAI, days after imbibition; DAP, days after pollination; LR, lateral roots; ME, malic enzyme; MS, Murashige and Skoog medium; PR, primary root; qPCR, real time polymerase chain reaction assay; RH, relative humidity; U, unit; WT, wild type; YFP, yellow fluorescent protein.

## Acknowledgements

CSA, MFD, MCGW and MS belong to the Researcher Career of National Council of Scientific and Technical Research (CONICET); CLA, TP, MBB and MG are fellows of the same institution. This work has been financially supported by National Agency for Promotion of Science and Technology, CONICET and Deutsche Forschungsgemeinschaft. The authors thank Rodrigo Vena, María J. Maymó, Marcos A. Tronconi, María C. Craia and Luisina Vitor Horen for the technical assistance to carry out this work.

## Author Contributions Statement

CLA, TP, GT, MBB, MG, MCGW and MS performed the experiments. VGM contributed to the plant lines. CLA, CSA, MFD, MCGW and MS planned the experiments, analyzed the data, and wrote the paper.

---

**Supplementary Figure 1:**
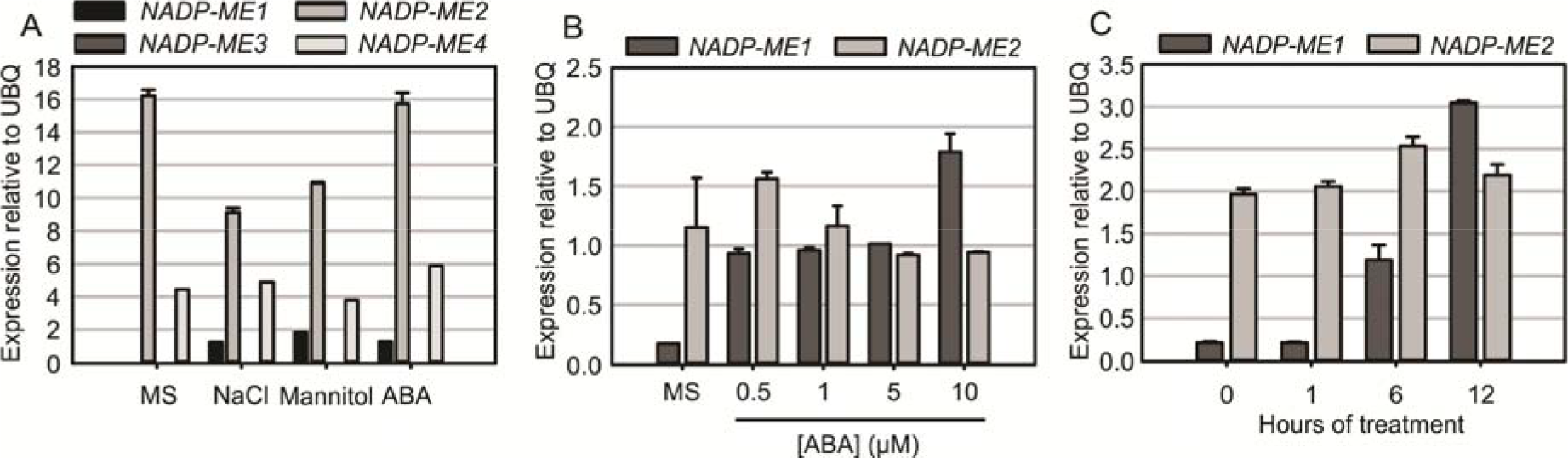
NADP-ME1 expression in response to NaCl, mannitol and ABA. (A) Relative levels of the transcripts of NADP-ME genes in rosettes in control conditions (MS) or after 6 hs treatments of seedling with 100 mM NaCl, 225 mM mannitol or 10 μM ABA. The expression levels of NADP-ME genes are normalized to expression of the reference gene polyubiquitin 10 (UBQ). The values are the average of at least two independent experiments ± SD. In (b) and (C) the levels of NADP-ME1 and NADP-ME2 transcripts after 12 h of ABA treatment or 10 μM ABA, respectively, are shown.

**Table S1: Motifs selected for the analysis of occurrence in promoters and 5’UTR.**

**Table S2: Location of motifs in the promoters and 5’UTR of the genes selected.**

**Table S3: Location of motifs in the promoters and 5’UTR of NADP-ME genes from *P. patens*.**

